# Macrophage- and CD4^+^ T cell-derived SIV differ in glycosylation, infectivity and neutralization sensitivity

**DOI:** 10.1101/2024.01.02.572735

**Authors:** Christina B. Karsten, Falk F.R. Buettner, Samanta Cajic, Inga Nehlmeier, Berit Roshani, Antonina Klippert, Ulrike Sauermann, Nicole Stolte-Leeb, Udo Reichl, Rita Gerardy-Schahn, Erdmann Rapp, Christiane Stahl-Hennig, Stefan Pöhlmann

**Affiliations:** Institute for the Research on HIV and AIDS-associated Diseases, University Hospital Essen, University of Duisburg-Essen, 45147 Essen, Germany; Institute of Clinical Biochemistry, Hannover Medical School, 30625 Hannover, Germany; glyXera GmbH, 39120 Magdeburg, Germany; Bioprocess Engineering Group, Max Planck Institute for Dynamics of Complex Technical Systems, 39106 Magdeburg, Germany; Infection Biology Unit, German Primate Center – Leibniz Institute for Primate Research, 37077 Göttingen, Germany; Unit of Infection Models, German Primate Center – Leibniz Institute for Primate Research, 37077 Göttingen, Germany; Nuvisan ICB GmbH, 13353 Berlin, Germany; Faculty of Biology and Psychology, Georg-August-University Göttingen, 37073 Göttingen, Germany

**Keywords:** infectivity, macrophage, neutralization, SIV, CD4^+^ T cell

## Abstract

The human immunodeficiency virus (HIV) envelope protein (Env) mediates viral entry into host cells and is the primary target for the humoral immune response. Env is extensively glycosylated, and these glycans shield underlying epitopes from neutralizing antibodies. The glycosylation of Env is influenced by the type of host cell in which the virus is produced. Thus, HIV is distinctly glycosylated by CD4^+^ T cells, the major target cells, and macrophages. However, the specific differences in glycosylation between viruses produced in these cell types have not been explored at the molecular level. Moreover, the impact of these differences on viral spread and neutralization sensitivity remains largely unknown. To address these questions, we employed the simian immunodeficiency virus (SIV) model. Glycan analysis revealed higher relative levels of oligomannose-type *N*-glycans in SIV from CD4^+^ T cells (T-SIV) compared to SIV from macrophages (M-SIV), and the complex-type *N*-glycans profiles differed between the two viruses. Notably, M-SIV demonstrated greater infectivity than T-SIV, even when accounting for Env incorporation, suggesting that host cell-dependent factors influence infectivity. Further, M-SIV was more efficiently disseminated by HIV binding cellular lectins. We also evaluated the influence of cell type-dependent differences on SIV’s vulnerability to carbohydrate binding agents (CBAs) and neutralizing antibodies. T-SIV demonstrated greater susceptibility to mannose-specific CBAs, possibly due to its elevated expression of oligomannose-type *N*-glycans. In contrast, M-SIV exhibited higher susceptibility to neutralizing sera in comparison to T-SIV. These findings underscore the importance of host cell-dependent attributes of SIV, such as glycosylation, in shaping both infectivity and the potential effectiveness of intervention strategies.

## Introduction

More than four decades since its discovery, the human immunodeficiency virus (HIV) and the associated disease, acquired immunodeficiency syndrome (AIDS), remain a significant global health challenge. In 2021, UNAIDS reported that 1.3 million individuals contracted HIV, and 630,000 AIDS-related deaths were observed (UNAIDS 2023). To combat this ongoing crisis, the development of vaccines and innovative antiviral strategies is crucial. The success of these initiatives will be based on a profound understanding of the complex interplay between HIV and its primary host cells, CD4^+^ T cells and macrophages.

The viral envelope protein, Env, mediates entry of HIV into host cells and constitutes the sole target for neutralizing antibodies (Walsh & Seaman 2021). Env is synthesized as an inactive precursor protein, gp160, in the secretory pathway of infected cells. During its trafficking through the Golgi apparatus, gp160 is proteolytically cleaved by furin into the surface unit, gp120, and the transmembrane unit, gp41 (Hallenberger et al. 1992), which remain non-covalently associated. For host cell entry, gp120 binds to the CD4 receptor and a chemokine coreceptor, usually C-C motif chemokine receptor 5 (CCR5) and/or C-X-C motif chemokine receptor 4 (CXCR4). Binding to receptor and coreceptor activates gp41, which drives the fusion of the viral and the target cell membranes, enabling the delivery of the viral genetic information into the host cell cytoplasm (Chen 2019).

A hallmark of Env, a trimeric type I transmembrane protein, is its extensive glycosylation, particularly of gp120, accounting for roughly 50 % of the molecular mass (Zhu et al. 2000). The glycans play a key role in viral spread: they shield underlying epitopes from attack by neutralizing antibodies (Wei et al. 2003) and facilitate viral capture by immune cell lectins like dendritic cell-specific intercellular adhesion molecule-grabbing nonintegrin (DC-SIGN) (Geijtenbeek et al. 2000), which likely play an important role in mucosal transmission. The fundamental importance of the glycan shield is underscored by its adaptation in response to the humoral immune response (Wei et al. 2003), and disrupting *N*-glycosylation signals of Env can render HIV (Koch et al. 2003; Ma et al. 2011; Quinones-Kochs et al. 2002) and the closely related simian immunodeficiency virus (SIV) susceptible to neutralization (Johnson et al. 2003; Reitter et al. 1998).

The process of *N*-glycosylation involves the transfer of a preformed oligosaccharide precursor linked to dolichol-phosphate (Dol-P-P-GlcNAc_2_Man_9_Glc_3_) onto asparagine residues within the consensus sequon Asn-X-Ser/Thr of nascent Env by the ER-localized oligosaccharyltransferase (Stanley et al. 2022). Following precursor attachment, initial trimming steps carried out by highly conserved ER resident glycosidases occur, which together with the action of glycosyltransferase, play pivotal roles in the regulation of Env folding and transport (Stanley et al. 2022). Only the fully folded trimeric Env enters the Golgi apparatus, where oligomannose-type glycans, generated in the ER, undergo further trimming and extension into hybrid and complex forms, potentially containing fucose, galactose, *N*-acetylglucosamine (GlcNAc), and sialic acid (Stanley et al. 2022). However, the level of *N*-glycan processing depends on the quaternary structure of Env and more than half of the *N*-glycans are not fully accessible to enzymatic processing due to their recessed location or to extremely dense glycan packaging, and thus remain in the oligomannose-type state (Pritchard et al. 2015; Zhu et al. 2000).

Notably, glycosylation of Env and cellular proteins is a cell type-dependent process (Liedtke et al. 1994; Liedtke et al. 1997; Raska et al. 2010; Willey et al. 1996) and differences in *N*-glycosylation of HIV and SIV from macrophages and CD4^+^ T cells have been associated with differential infectivity (Gaskill et al. 2008; Heeregrave et al. 2023), neutralization sensitivity (Heeregrave et al. 2023; Willey et al. 1996) and lectin reactivity (Heeregrave et al. 2023; Lin et al. 2003). Despite their importance in virus-host cell interactions and immune control, host cell-specific glycosylation differences have not been determined for HIV or SIV at a molecular level. Furthermore, a detailed comparative analysis of the biological properties of isogenic HIV and SIV produced in macrophages and CD4^+^ T cells has been lacking. In this study, we aim to address these knowledge gaps by strategically investigating the influence of macrophage or CD4^+^ T cell origin on the significance of Env glycosylation, viral spread, and neutralization sensitivity, using SIV as a model for HIV.

## Results

### Production of SIV in macrophages and CD4^+^ T cells

To produce isogenic SIV in CD4^+^ T cells and macrophages, it was imperative to employ a molecularly cloned SIV variant capable of robust replication in both cell types. Our choice for this purpose was a SIVmac239 variant, specifically SIVmac239/316 Env, characterized by nine amino acid substitutions within Env in comparison to the parental strain (Mori et al. 1993; Mori et al. 1992). These substitutions facilitate efficient utilization of CCR5 in the absence of or at very low levels of CD4 expression (Puffer et al. 2002), a condition observed in rhesus macaque macrophages (Mori et al. 2000). For the preparation of SIVmac239/316 Env stocks, CD4^+^ T cells and macrophages were generated from rhesus macaque peripheral blood mononuclear cells (PBMCs) and their identity was confirmed by analysis of cell surface marker expression via flow cytometry (Fig. S1).

Subsequently, CD4^+^ T cell and macrophage cultures were infected with SIVmac239/316 Env, input virus was removed, and the culture supernatants harvested over a two-week period. Notably, pooled viral supernatants from CD4^+^ T cells (T-SIV) contained 6.5-fold more p27-capsid antigen per ml on average than viral supernatants from macrophages (M-SIV), as determined by enzyme-linked immunosorbent assay (ELISA) (data not shown). Finally, a comprehensive assessment of the complete *env* sequences, which were amplified through reverse transcriptase-polymerase chain reaction (RT-PCR) from the culture supernatants, provided confirmation that no genetic mutations were introduced throughout the *in vitro* propagation process.

### The quantity of oligomannose-type and certain complex *N*-glycans of gp120 differs between M-SIV and T-SIV

The glycan shield of both HIV and SIV plays a pivotal role in immune evasion and mediating lectin-dependent interactions with immune cells. It is plausible that these functions could be modulated by cell type-specific alterations in the glycosylation pattern. To investigate this, we subjected concentrated virions to enzymatic digestion using endoglycosidase H (Endo H), which selectively removes oligomannose-type and certain hybrid *N*-glycans, and peptide-*N*-glycosidase F (PNGase F), which eliminates all *N*-linked glycans. The band shift upon Endo H digestion was more pronounced for T-SIV than M-SIV suggesting that T-SIV gp120 is adorned with a higher proportion of oligomannose-type glycans compared to M-SIV gp120 (Fig. 1). These findings align with previously published data (Gaskill et al. 2008; Lin et al. 2003; Willey et al. 1996) but do not offer insights into the specific structures of the differentially incorporated *N*-glycan species.

**Fig. 1.**
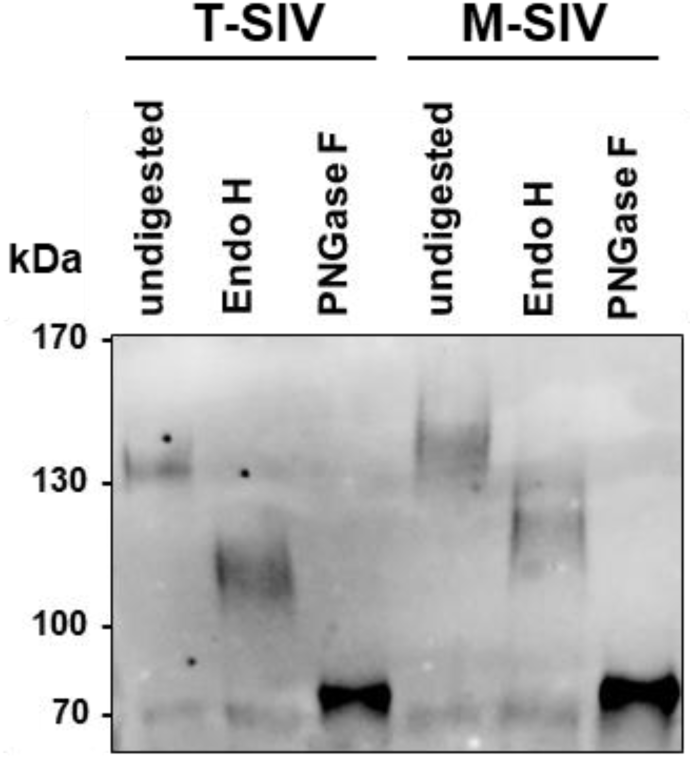
CD4^+^ T cell-derived simian immunodeficiency virus (T-SIV) gp120 carries more oligomannose-type glycans than gp120 of macrophage-derived SIV (M-SIV). A) T-SIV and M-SIV viral stocks were normalized for comparable gp120 content and concentrated. The viruses were subjected to mock treatment or enzymatic digestion with endoglycosidase H (Endo H) or peptide-*N*-glycosidase F (PNGase F), followed by western blot detection of the envelope protein (Env). Consistent results were obtained across three independent experiments.

To address the latter question, we performed *N*-glycan analytics of T-SIV and M-SIV gp120 by multiplexed capillary gel electrophoresis with laser-induced fluorescence detection (xCGE-LIF). Further, the identity of the proteins subjected to *N*-glycan analytics was confirmed by liquid chromatography-tandem mass spectrometry (LC-MS/MS) (data not shown). The xCGE-LIF fingerprints of *N*-glycans derived from T-SIV and M-SIV gp120 show considerable differences in the relative intensities of the detected peaks suggesting quantitative differences in the incorporated glycan species (Fig. 2A, supplementary table 1).

**Fig. 2.**
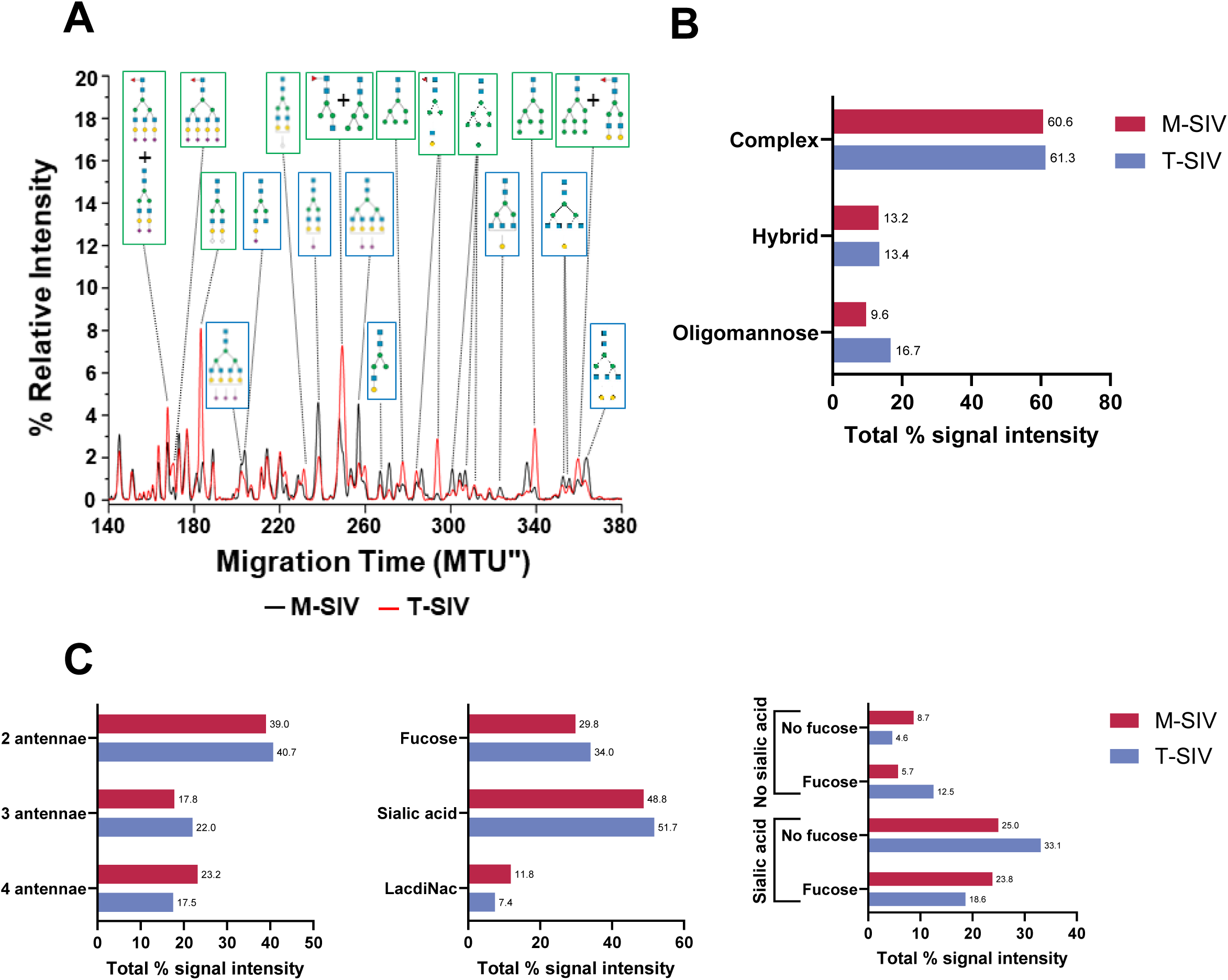
Differences in the relative distribution of *N*-glycans from gp120 of M-SIV and T-SIV measured by multiplexed capillary gel electrophoresis with laser-induced fluorescence detection (xCGE-LIF). A) The electropherogram region with the most striking differences between M-SIV and T-SIV is plotted (140-380 migration time units (MTU”)). Peak intensities are presented as percentage of the total peak height to obtain the relative signal intensity (in %) for each peak (representing at least one *N*-glycan structure). A selection of distinct *N*-glycan structures enriched in T-SIV are denoted with green boxes, while those enriched in M-SIV are marked with blue boxes. Symbolic representation of *N*-glycan structures in the figure were drawn with the software GlycanBuilder2 (Tsuchiya et al. 2017), in alignment with the updated guidelines of the Symbol Nomenclature For Glycans working group (Neelamegham et al. 2019): green circle, mannose; yellow circle, galactose; blue square, *N*-acetylglucosamine; pink diamond, *N*-acetylneuraminic acid; white diamond, *N*-glycolylneuraminic acid; red triangle, fucose. B, C) To delve deeper into the distinctions in the *N*-linked glycan profile of M-SIV and T-SIV gp120, we aggregated the xCGE-LIF signal intensities of peaks corresponding to glycan species associated with specific groups to calculate the total % signal intensity. These data were calculated based on the information provided in supplementary table 1. B) illustrates the distribution of annotated glycan species across various glycan types, while C) categorizes complex glycans based on their distinct features. Numbers at the end of bars give the exact total % signal intensity for the specific glycan group.

A relative quantification of *N*-glycan signal intensities was performed. For this, the total peak intensity (tpi) of a peak was allocated to all *N*-glycan groups of the analysis, for which the assigned single or multiple glycan species met the criteria. This analysis revealed the following: T-SIV exhibited increased relative levels of oligomannose structures on gp120 compared to M-SIV (Fig. 2B, 16.7 % vs. 9.6 % of the tpi); in alignment with the results from glycosidase digestion and subsequent western blot analysis (Fig. 1). Additionally, profiles of complex-type *N*-glycans differed between the two viruses (Fig. 2C). M-SIV gp120 displayed more extensive branching of complex glycans, with a higher proportion featuring four antennae rather than three, in comparison to T-SIV (23.2 % vs. 17.5 % tpi). An overall assessment of fucose, sialic acid, and LacdiNAc content revealed that T-SIV gp120 harbors increased levels of *N*-glycan species carrying fucose (34 vs. 29.8 tpi) and sialic acid (51.7 vs. 48.8 tpi) compared to M-SIV; while the latter exhibited more LacdiNAc (11.8 vs. 7.4 tpi) containing structures. A more detailed analysis of the complex -type glycan species revealed a more nuanced pattern: M-SIV complex glycans tended to feature either both fucose and sialic acid residues (23.8 (M-SIV) vs. 18.6 % tpi (T-SIV)) or neither (8.7 (M-SIV) vs. 4.6 % tpi (T-SIV)). Differently, T-SIV complex glycans tended to have either fucose (12.5 (T-SIV) vs. 5.7 % tpi (M-SIV)) or sialic acid (33.1 (T-SIV) vs. 25 % tpi (M-SIV)) but not both. Taken together, despite general similarities in the *N*-glycomes of M-SIV and T-SIV gp120, there are pronounced differences in the level of oligomannose-type *N*-glycans and their complex *N*-glycan profiles.

### M-SIV is better equipped for both direct and indirect viral spread compared to T-SIV

The revelation that T-SIV and M-SIV exhibit significant disparities in glycan coat composition raised the question of whether these distinctions are linked to variations in other SIV characteristics pertinent to viral dissemination. To investigate this possibility, our initial focus was on viral infectivity. When TZM-bl indicator cells were exposed to viruses normalized for p27-capsid content, it became evident that M-SIV exhibited significantly higher infectivity than T-SIV (Fig. 3A, 2-way ANOVA, p = 0.03). However, both viruses exhibited *in vivo* infectivity and replicated to comparable levels (Fig. S2), and when we equalized the infectivity of T-SIV and M-SIV stocks determined on C8166 T cells, the infection efficiency was found to be similar (Fig. 3B). These results underscore that the infectivity of M-SIV per ng capsid antigen surpasses that of T-SIV, confirming the findings previously obtained for SIV (Gaskill et al. 2008) and dual-tropic HIV (Heeregrave et al. 2023).

**Fig. 3.**
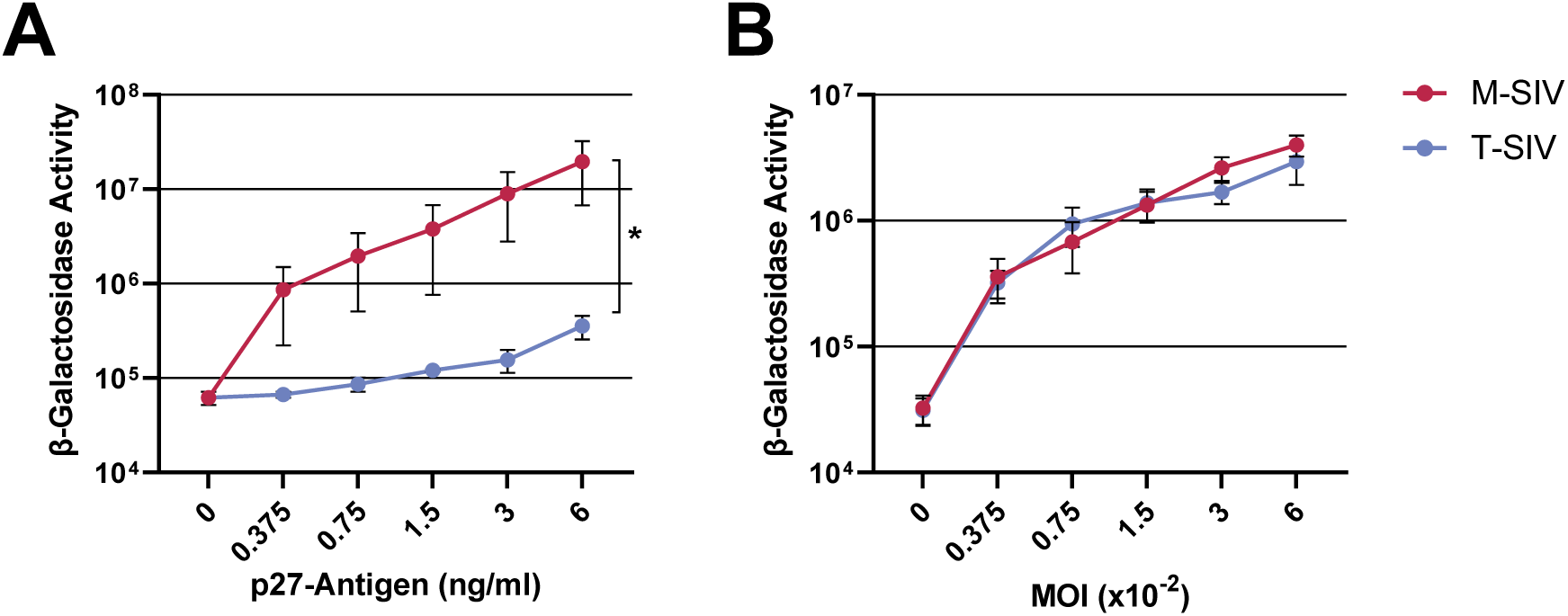
M-SIV is more infectious than T-SIV. A) TZBM-bl indicator cells were exposed to equal volumes of M-SIV and T-SIV stocks, normalized for p27-capsid protein (left panel) or infectious units (right panel). Following virus removal at 5-6 hours post-infection, β-galactosidase activity was measured in cell lysates 72 hours post-infection. The grand mean of three experiments performed in triplicates (left panel) or quadruplicates (right panel) are shown with the standard error of the mean (SEM). Statistical significance between datasets determined by two-way ANOVA (*, p ≤ 0.05).

Glycosylation also affects HIV Env incorporation into viral particles (Li et al. 1993). To clarify whether the increased M-SIV infectivity in comparison to T-SIV could be attributed solely to disparities in Env incorporation, we subjected p27-capsid normalized quantities of both M-SIV and T-SIV to western blot analysis to assess their gp120 and p27 content, and quantified the signal using the software ImageJ (Fig. 4). A visual examination already revealed that the less infectious T-SIV exhibited lower levels of gp120 incorporation compared to the more infectious M-SIV (Fig. 4A). Following quantification and subsequent normalization of the gp120 signal to the p27-capsid signal, the data indicated a noteworthy 62 % reduction in Env incorporation for T-SIV when compared to M-SIV (Fig. 4B, t-test, p = 0.0051). While this effect is significant, the disparities in infectivity between the two viruses (55-fold at 6 ng p27/ml) surpasses the differences in Env incorporation. This suggests that the virus-producing cell has a broader impact on SIV infectivity beyond its influence on Env incorporation.

**Fig. 4.**
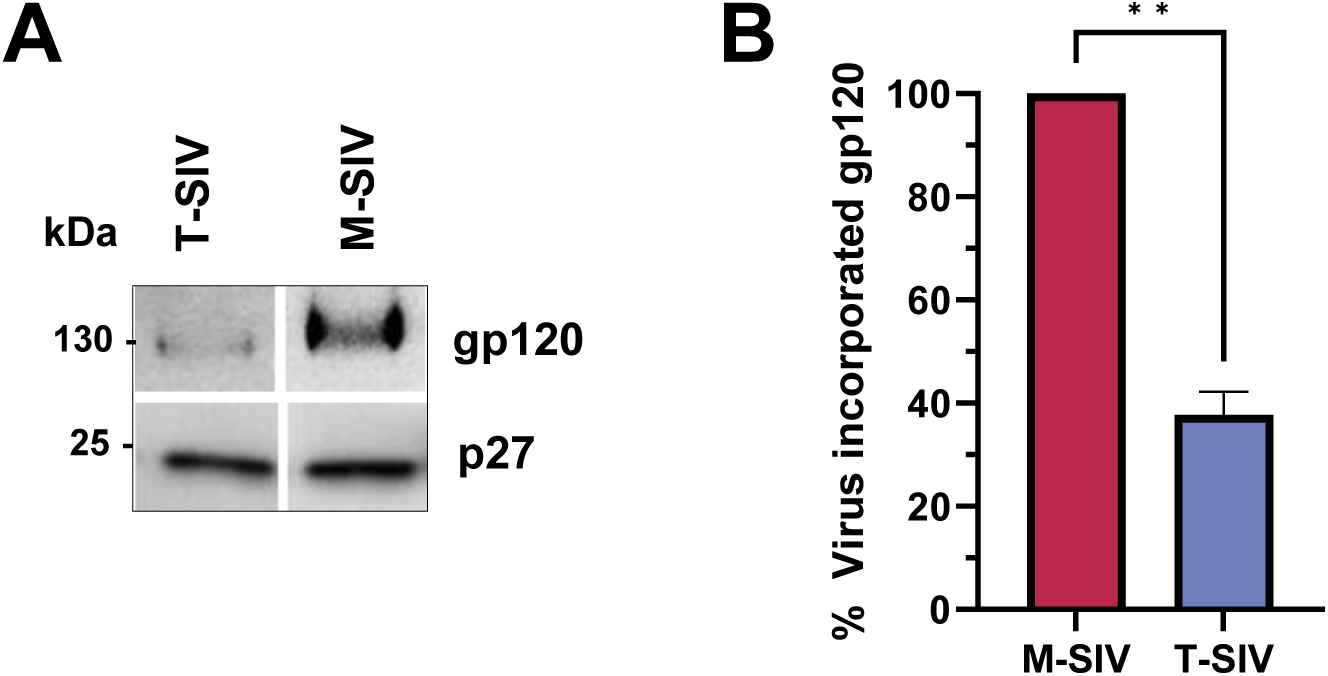
M-SIV incorporates more gp120 than T-SIV. A) M-SIV and T-SIV normalized for equal amounts of p27 were concentrated, resolved using SDS-PAGE, and analyzed by western blot for gp120 and p27. Consistent outcomes were observed across three independent experiments. B) The software ImageJ (Schneider et al. 2012) was utilized to quantify gp120 and p27 signal intensities obtained in A). The gp120 signal per p27 signal ratio was calculated, and values were plotted, with M-SIV set as 100 %. Presented is the grand mean with SEM. Paired t-test was applied to assess statistical differences between groups (**, p ≤ 0.01).

Lectins modulate HIV and SIV transmission to susceptible cells *in vitro* (Bashirova et al. 2001; de Witte et al. 2007; Geijtenbeek et al. 2000; Pöhlmann et al. 2001), a process possibly relevant for early viral spread after transmission *in vivo* (Gonzalez et al. 2019). Since both M-SIV and T-SIV were found to be infectious in rhesus macaques via the rectal route (Fig. S2), the question arose whether the cell type-dependent differences in these viruses influence their engagement of lectin receptors. Specifically, the interactions of M-SIV and T-SIV with the lectins with potential positive (DC-SIGN (Geijtenbeek et al. 2000), DC-SIGN related protein (DC-SIGNR) (Bashirova et al. 2001)) or negative (Langerin (de Witte et al. 2007)) influence on HIV/SIV viral spread were investigated. For this purpose CEMx174 R5 target cells were infected in two ways: directly with either virus, normalized for equal infectivity, or indirectly via co-culture with Raji transmitter cells bearing either no lectin or the aforementioned lectins (Fig. 5). While direct infection of target cells confirmed the equivalent infectivity of M-SIV and T-SIV, and co-culturing the virus with transmitter cells in the absence of target cells yielded only background signals, a significant difference emerged. M-SIV was more efficiently transferred to target cells by DC-SIGN compared to T-SIV (t-test, p = 0.01). This observation is consistent with previous findings related to the transmission of dual-tropic HIV produced in macrophages versus CD4^+^ T cells through DC-SIGN (Heeregrave et al. 2023). Similarly, DC-SIGNR and Langerin preferentially transferred M-SIV in comparison to T-SIV (t-test, p = 0.0001, 0.0003). These findings suggest that SIV replication in macrophages produces viral particles with distinctive characteristics that confer a clear advantage in direct and possibly indirect viral spread via lectins compared to viruses originating from CD4^+^ T cells.

**Fig. 5.**
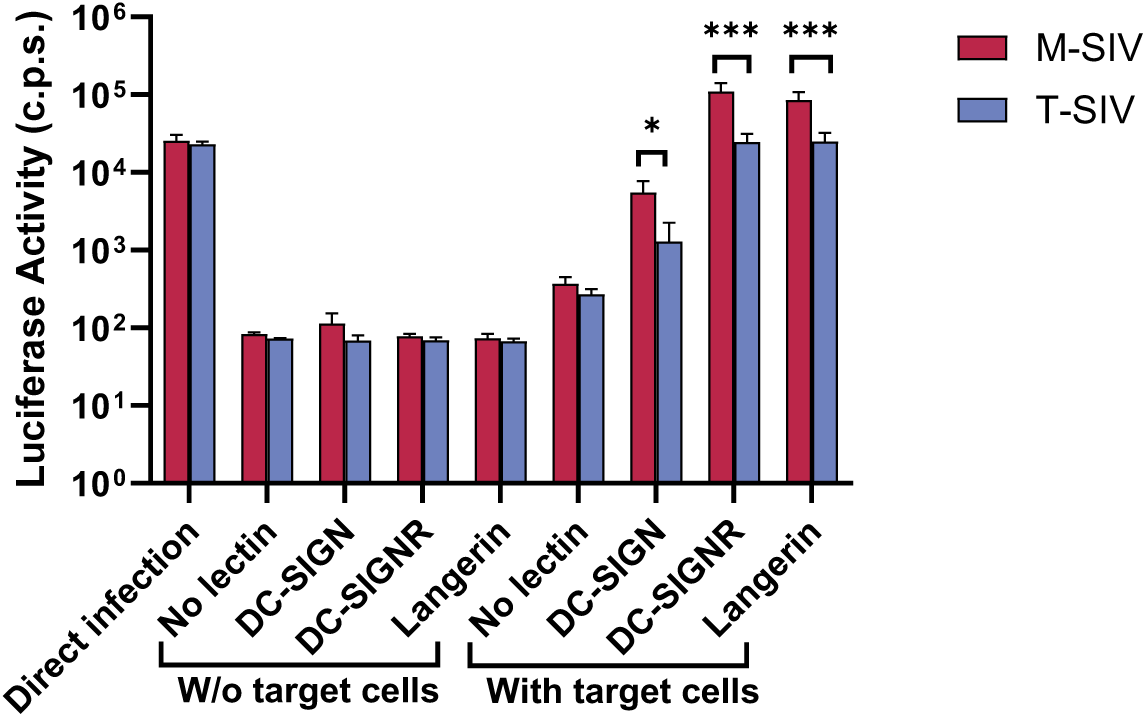
Lectins transmit M-SIV better than T-SIV. M-SIV and T-SIV were incubated with Raji cells expressing no additional lectin, dendritic cell-specific intercellular adhesion molecule-grabbing nonintegrin (DC-SIGN), DC-SIGN related protein (DC-SIGNR) or Langerin. Unbound virus was removed and the transmitter cells were co-cultured with CEMx174 R5 target cells. Infection of target cells was detected by the measurement of luciferase activity in cell lysates 72 h post infection. Direct infection of CEMx174 R5 cells served as positive control, while negative controls consisted of Raji cells incubated without target cells. The grand mean of three independent experiments conducted in triplicates with SEM, normalized to direct target cell infection, is depicted. Statistical differences between M-SIV and T-SIV for each transmitting lectin were calculated by t-test (*, p <= 0.05; ***, p <= 0.001 ). C.p.s.: counts per second.

### Host cell-dependent features of SIV define the sensitivity of M-SIV and T-SIV to carbohydrate binding agents (CBAs) and neutralizing antibodies

CBAs and especially broadly neutralizing antibodies are under investigation as promising biomedical treatments to prevent the spread of HIV (Julg & Barouch 2021; Nabi-Afjadi et al. 2022). They ultimately target Env and must interact with or overcome its dense glycan shield. Therefore, it was plausible that host cell-derived features of SIV Env, such as the observed differences in glycomes between M-SIV and T-SIV, might influence the effectiveness of CBAs and neutralizing antibodies. To explore this hypothesis, we examined the ability of CBAs to inhibit SIV infection of TZM-bl cells, focusing on two glycan species groups: oligomannose glycans and core fucosylated complex glycans, which were incorporated in different quantities into M-SIV and T-SIV gp120 (Fig. 2). Pre-treatment with ulex europaeus agglutinin (UEA), a lectin against core fucose, did not interfere with the infection of TZM-bl cells by controls (HIV-1 or vesicular stomatitis virus glycoprotein (VSV-G) pseudotyped viruses) or SIV (Fig. 6A) as expected (Karsten et al. 2015). As anticipated, HIV-1 was highly sensitive to inhibition by mannose-specific lectins cyanovirin-N (CV-N, target: Manα1-2) or galanthus nivalis agglutinin (GNA, target: Manα1-3, Manα1-6), while there was significantly less inhibition of infection by pseudotypes bearing the minimally glycosylated protein VSV-G (Balzarini et al. 1991; Boyd et al. 1997). Consistent with the xCGE-LIF data (Fig. 2), T-SIV with its higher oligomannose content in gp120 was more strongly inhibited than M-SIV in a concentration-dependent manner (Fig. 6A), and these differences were statistically significant (Fig. 6B, t-test, CV-N: p = 0.0032, GNA: p = 0.008). Thus, the producer cell type can influence SIV susceptibility to inhibition by mannose-specific CBAs.

**FIG 6.**
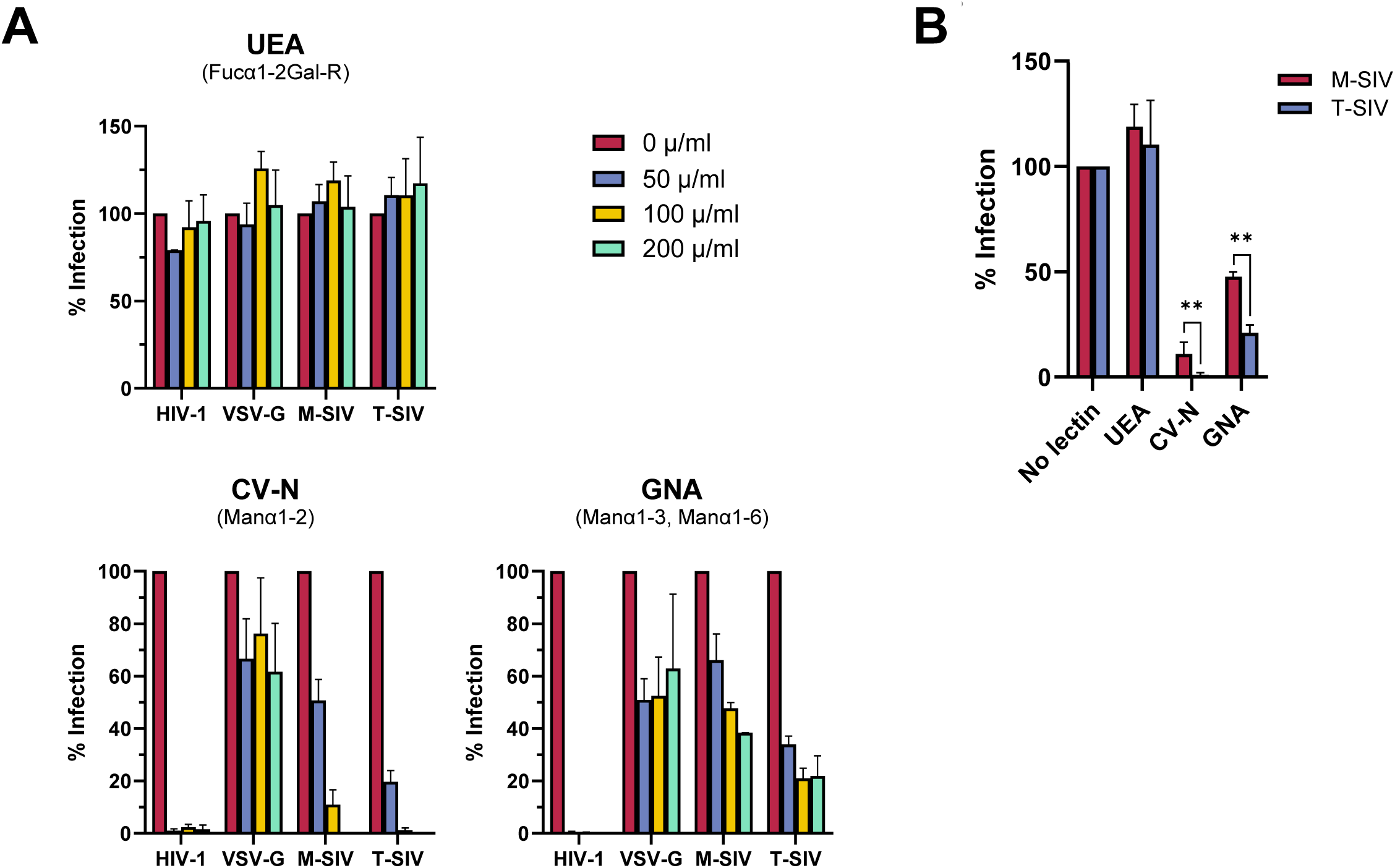
T-SIV is more sensitive to inhibition by mannose-specific lectins than M-SIV. Infectivity-normalized M-SIV, T-SIV, HIV-1 NL4-3 and *env*-defective HIV-1 NL4-3 pseudotyped with vesicular stomatitis virus glycoprotein (VSV-G) were preincubated with the indicated concentrations of ulex europaeus agglutinin (UEA), cyanovirin-N (CV-N), or galanthus nivalis agglutinin (GNA), before addition to TZM-bl indicator cells. Virus was removed at 5-6 h post infection and β–galactosidase activity was measured in cell lysates at 72 h post infection. Presented are the grand mean values with SEM normalized to lectin-free conditions from two independent experiments conducted in triplicates for all lectin concentrations. B) Plotted are the results obtained in A) for M-SIV and T-SIV using a lectin concentration of 100 µg/ml. Statistical differences between M-SIV and T-SIV were calculated for each lectin using a paired t-test (**, p ≤ 0.01).

To investigate whether cell type-dependent differences in SIV impact antibody neutralization effectiveness, we conducted infections of TZM-bl cells with HIV, M-SIV, and T-SIV in the presence of neutralizing sera. These sera were obtained from rhesus macaques infected with SIVmac239, the parental strain of SIVmac239/316 Env used for producing M- and T-SIV. This parental virus differs in nine amino acids, making it more resistant to antibody neutralization (Puffer et al. 2002). Using four different sera, we observed some inhibition of HIV (Fig. 7A). Intriguingly, M-SIV was neutralized more efficiently than T-SIV. These effects were concentration-dependent (Fig. 7B) and statistically significant (Fig. 7C, paired t-test, p = 0.013). In light of these findings, we conclude that differences originating from the virus-producing cell can influence the efficacy of CBA and antibody neutralization, and thus might have relevance for the development of biomedical interventions based on these scaffolds.

**FIG 7.**
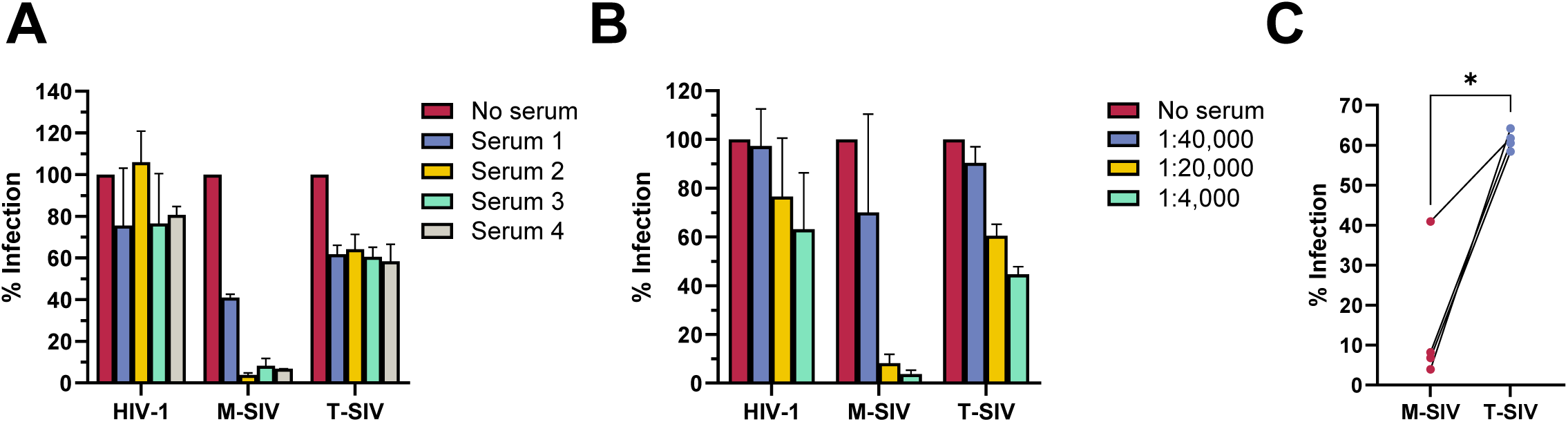
M-SIV is more sensitive to serum neutralization than T-SIV. A) Infectivity-normalized M-SIV, T-SIV, and HIV-1 NL4-3 were preincubated with sera from SIVmac239-infected rhesus macaques at varying dilutions prior to infection of TZM-bl indicator cells. Incubation of virus with medium alone served as negative control. Virus removal occurred at 5-6 hours post-infection, with β-galactosidase activity measured in cell lysates at 72 hours post-infection. Shown is the grand mean with SEM from two independent experiments conducted in triplicates at a serum dilution of 1:20,000. Infection in the absence of serum was normalized to 100 %. B) The titration curve for serum 3 from A) is presented as grand mean with SEM. C) Data for sera 1-4 from A), all at a 1:20,000 dilution, are directly compared. Paired t-test was applied to assess differences in means between groups (*, p ≤ 0.05).

## Discussion

Here, we analyzed whether an infectious molecular clone of SIV produced in primary rhesus macaque macrophages (M-SIV) or CD4^+^ T cells (T-SIV) differs in Env glycosylation, viral spread and neutralization sensitivity. We found that the overall glycan landscape of M-SIV and T-SIV gp120 shared similarities; however, different quantities of the glycan species were incorporated. While oligomannose-type *N*-glycans were more frequent in T-SIV gp120, the complex glycan profiles between both viruses varied considerably. The producer cell type also influenced virus characteristics, which are important for viral spread with M-SIV being more infectious, incorporating more Env, and being better transmitted by lectin receptors than T-SIV. We further found host cell-dependent differences in viral sensitivity to CBAs and neutralizing sera showing that T-SIV, bearing higher levels of oligomannose structures on gp120, was more sensitive to mannose-specific lectins, while M-SIV was more efficiently neutralized by sera.

It has been documented previously that CD4^+^ T cell-derived viruses exhibit a greater prevalence of oligomannose structures in gp120 (Gaskill et al. 2008; Lin et al. 2003) but less LacDiNac residues in comparison to virus originating from macrophages (Willey et al. 1996). By providing the first comparative molecular analysis of gp120 glycosylation of primary host cells, we corroborate and expand upon these earlier observations. Our xCGE-LIF analysis demonstrated an increased presence of M5, M6, and M8 on T-SIV-derived gp120 in comparison to M-SIV-derived gp120. The signature for M9 also appeared to be elevated in T-SIV versus M-SIV. However, it is important to note that this study did not allow for a definitive conclusion due to the potential co-migration of Man9 with another *N*-glycan structure (FA2G2). Nevertheless, other studies have indicated that Man9 is a frequently incorporated glycan species, or even the predominant one, in CD4^+^ T cell-derived HIV gp120 (Bonomelli et al. 2011; Panico et al. 2016). The disparities in the complex glycan profiles of M-SIV and T-SIV gp120 reveal that variations in the incorporation of LacDiNac are just one aspect of the distinct glycomes originating from these two cell types. These distinctions align with the expectations generated by an mRNA analysis of more than 20 glycosylation-related enzymes, which exhibit differential expression in CD4^+^ T cells and macrophages (Gaskill et al. 2008). Specifically, rhesus macaque CD4^+^ T cells exhibit elevated expression levels of genes that potentially enhance core fucosylation (FUT8) and sialylation (ST6Gal1), while rhesus macaque macrophages express higher levels of genes that promote the conversion of oligomannose to complex glycans (MGAT1), a reduction in sialic acid (NPL), and high-level branching (MGAT4B) (Gaskill et al., 2008). Finally, our findings indicate that M-SIV gp120 displays a more extensive branching of complex glycans, a higher frequency of glycan species featuring both core fucose and sialylation, and a greater LacDiNac content compared to T-SIV gp120. This suggests that gp120 might undergo a higher degree of glycan processing in macrophages than in CD4^+^ T cells.

The advantages observed for M-SIV over T-SIV in terms of direct and indirect viral spread prompt the question of whether these differences could lead to more efficient *in vivo* transmission of macrophage-derived viruses. The characteristics of successfully sexually transmitted HIV isolates are, apart from being usually R5-(Kariuki et al. 2017) and likely T cell-tropic (Alexander et al. 2010), under debate. Interestingly, recent studies demonstrated direct infection of macrophages with HIV by fusion with infected CD4^+^ T cells (Han et al. 2022) and indirectly by infection with CD4^+^ T cell-derived HIV transcytosed through epithelial cells (Real et al. 2018). Further, macrophages were determined as an important viral reservoir in penile tissue (Ganor et al. 2019) and the main virus source in semen (Fennessey et al. 2022). In light of these recent findings, it is noteworthy that SIVmac239/316 Env produced in macrophages exhibited traits similar to those reported for HIV-1 clade B and C transmitted founder viruses when compared to unmatched chronic isolates. In this study, transmitted founder viruses showed increased Env incorporation, higher infectivity, and enhanced transfer to target cells by DCs in comparison to chronic viral isolates (Parrish et al. 2013). Similarly, SIVmac239/316 Env replicated to higher levels in CD4^+^ T cells as compared to macrophages and M-SIV incorporated more Env, was displaying higher infectivity, and was better transmitted by (DC) lectins than T-SIV. This suggests that macrophage origin might further strengthen viral characteristics, which have already been associated with successful viral transmission. Furthermore, one of our previous studies using SIVmac239/316 Env and the rhesus macaque model indicated that exclusive oligomannose glycosylation of Env completely prevents *in vivo* SIV transmission (Karsten et al., 2015), suggesting that the oligomannose glycan content of CD4^+^ T cell-derived HIV and SIV may be unfavorable during sexual transmission. While our exploratory animal experiment showed that the oligomannose profile of T-SIV Env did not have the same detrimental effect on virus transmission as observed for exclusive oligomannose glycosylation, larger animal studies might decipher potential differences in the *in vivo* transmissibility of macrophage and CD4^+^ T cell-derived viruses. Thus, transmitted founder viruses possibly replicate in macrophages prior and after transmission, and macrophage-dependent viral traits such as the Env glycosylation profile might booster virus dissemination.

Our results support the conclusions of others that cell type-dependent differences in HIV and SIV, like Env glycosylation, influence the sensitivity towards potential biological interventions including lectins and antibodies (Heeregrave et al. 2023; Raska et al. 2010; Willey et al. 1996). In contrast to our results, two other studies determined macrophage-derived HIV to be more neutralization sensitive than viruses produced in CD4^+^ T cells (Heeregrave et al. 2023; Willey et al. 1996). While one study utilized HIV and 2G12, a mannose-only binding neutralizing antibody, for their investigations (Heeregrave et al. 2023), the other study made this conclusion by using sera of an HIV infected chimpanzee (Willey et al. 1996). This suggests that whether macrophage or CD4^+^ T cell-derived traits provide protection for the virus *in vivo*, is likely dependent on the specific antibody profile of the host. Host cell-dependent neutralization sensitivity was recently also shown to introduce a bias into the selection of neutralizing antibodies for HIV clinical trials (Cohen et al. 2018). The standard assay for the assessment of candidate neutralizing antibodies utilizes 293T-derived pseudotypes with HIV envelopes of different strains (Sarzotti-Kelsoe et al. 2014), but the efficiency of antibody neutralization differed when the same viruses were produced in PBMCs instead (Cohen et al. 2018). The *in vitro* results suggest that cell host-dependent viral traits like Env glycosylation add another layer of complexity to Env diversity beyond genetic variability. Current efforts to design antibody-based therapy approaches to control genetically complex HIV swarms in infected individuals, aim to target multiple epitopes on Env at once including glycan-protein targets (Wagh & Seaman 2023). Although host cell-dependent differences appear to be important for antibody and lectin interactions *in vitro*, these might have a negligible relevance *in vivo*, considering that CD4^+^ T cells are likely the main virus producing cells over the course of infection (DiNapoli et al. 2016). Future research must determine whether host cell-dependent viral distinctions should factor into the selection of candidate therapeutics and the design of strategies to address the challenges posed by the extensive diversity of HIV.

One limitation of this study is the use of a single SIV strain, which was carefully chosen from the few available macrophage-tropic SIV proviruses previously used in studies with rhesus macaques. Since the completion of our study, HIV strains were found to differ in their sensitivity to host cell-derived modifications of their traits (Heeregrave et al. 2023), emphasizing the need for testing multiple viral strains for more comprehensive conclusions. Additionally, limited blood availability from rhesus macaques constrained the production of sufficient virus for extended glycan analysis, preventing the identification of overlapping peaks in xCGE-LIF annotations via glycosidase digest. Finally, host cell-dependent glycosylation differences in Env have been repeatedly linked to observed variations in viral functions (Gaskill et al. 2008; Heeregrave et al. 2023; Lin et al. 2003). However, other factors such as viral surface glycosylation beyond Env (Spillings et al. 2022), host cell protein incorporation (Lawn et al. 2000; Munoz et al. 2022), and virus stock impurities like exosomes (Hazrati et al. 2022) could have influenced the presented results. Establishing a direct link between Env glycosylation and viral functions is technically challenging and beyond the scope of this study. Nevertheless, infectivity experiments with SIV after glycosidase treatment of the virus performed by Gaskill and colleagues, support a direct relationship between Env glycosylation and SIV infectivity (Gaskill et al. 2008).

In summary, our study using SIV as a model for HIV provides the first detailed molecular characterization of gp120 *N*-glycomes as derived by the target cells macrophages and CD4^+^ T cells. Further, it highlights the significance of host cell-dependent differences *in vitro*, affecting both direct and indirect viral spread, as well as neutralization sensitivity. Overall, our findings might have broader implications for the successful development of innovative strategies for HIV prevention and therapy.

## Material and Methods

### Animal studies

The animal studies were conducted at the German Primate Center, which has the permission to breed and house non-human primates under license number 392001/7 granted by the local veterinary office and conforming with x 11 of the German Animal Welfare act. Ethics approval was obtained from a committee authorized by the Lower Saxony State Office for Consumer Protection and Food Safety with the project license 33.14-42502-04-11/026. All animals were bred, cared for by qualified staff and, housed at the German Primate Center adhering to the German Animal Welfare Act and complying with the European Union guidelines for the use of nonhuman primates in biomedical research.

In total, eight male and one female Indian-origin rhesus macaques (*Macaca mulatta*) were assigned to experimental groups based on their age (4 to 7 years). A maximum of 5-6 ml blood per kg bodyweight was drawn *ex vivo* from animals, which were anesthetized intramuscularly with 10 mg ketamine per kg body weight from the femoral vein employing the Vacutainer system (BD Biosciences). For virus challenges, animals were anesthetized intramuscularly with a mixture of 5 mg ketamine, 1 mg xylazin, and 0.01 mg atropine per kg body weight. Virus introduction occurred up to ten centimeters into the rectum using a catheter (Urotech). During the procedure and for the subsequent 30 minutes, the animals were maintained in a ventral position with an elevated pelvis. Monitoring for infection establishment by RT-PCR took place every two and three weeks post-challenge.

### Plasmids

The plasmids encoding HIV-1 NL4-3 (Pöhlmann et al. 2001), SIVmac239/316 Env (Mori et al. 1992), the HIV-1 NL4-3-derived vector pNL4-3.Luc.R-E- (Connor et al. 1995), and the vesicular stomatitis virus glycoprotein (Naldini et al. 1996) have been previously described.

### Cell culture

293T and TZM-bl cells were cultured in Dulbecco’s modified Eagle’s medium (DMEM; PAN-Biotech) supplemented with 10% fetal bovine serum (FBS, Biochrome), 100 U/ml penicillin, and 100 µg/ml streptomycin (P/S; PAN-Biotech). The suspension cell lines (C8166, CEMx174 R5, Raji, Raji DC-SIGN/DC-SIGNR/Langerin) were cultured in RPMI 1640 supplemented with L-glutamine (PAN-Biotech), 10 % FBS, and P/S. For the isolation of rhesus macaque primary CD4^+^ T cells and macrophages, PBMCs were isolated from whole blood using ficoll (Biochrom) density gradient centrifugation. CD4^+^ T cells were purified by negative depletion using magnetic beads (Miltenyi Biotech) and cultured in RPMI 1640 supplemented with 20 % FBS, P/S and 10 µg/ml concanavalin A (Sigma-Aldrich) for 24 h at a density of 2 x 10^6^ cells/ml. Following that, CD4^+^ T cells were cultured in RPMI 1640 supplemented with 20 % FBS, P/S and 100 U/ml recombinant human interleukin-2 (IL-2). For the generation of macrophages, monocytes were purified from PBMCs by positive selection for CD14^+^ cells with magnetic beads (Miltenyi Biotech). For differentiation into macrophages, monocytes were seeded at a density of 3 x 10^5^ cells/ml in RPMI 1640 medium supplemented with 20 % FBS, 10 % human AB serum (Sigma-Aldrich), and 10 ng/ml recombinant human macrophage colony stimulating factor (Peprotech) and cultured for 5 d. Subsequently, macrophages were cultured in RPMI 1640 supplemented with 20% FBS. All cells were grown in a humidified atmosphere at 37 °C with 5% CO_2_.

### Flow cytometry

For flow cytometric analysis of marker expression, 50,000 to 500,000 PBMCs and the above mentioned cell subsets were stained for 30 min at room temperature (RT) with mixtures of monoclonal antibodies (mAb) reactive against CD3 (SP34-2, Alexa Fluor 700), CD4 (L200, V450), CD11b (ICRF44, PE), CD16 (3G8, FITC) and CD20 (L27, PE-Cy7) all from BD Biosciences, as well as CD8 (3B5, Pacific Orange, Invitrogen), and CD14 (RMO52, ECD, Beckman Coulter) diluted in staining buffer (phosphate-buffered saline with 5% FBS). Subsequently, cells were washed with staining buffer, and fixed with 4 % paraformaldehyde solution for seven minutes. After an additional washing step, the cells were analyzed using a LSRII cytometer (BD Biosciences) equipped with three lasers. Compensation was calculated by FACS DIVA software 6.1.3 using appropriate single antibody labeled compensation beads from SpheroTech. Data analysis was performed using FlowJo software v9.6 (Treestar).

### Production of viruses and pseudotyped viruses

To generate virus stocks of HIV-1 NL4-3 and SIVmac239/316 Env, 293T cells were seeded into T25-cell culture flasks and transfected with 12 µg of plasmids encoding proviral DNA using calcium phosphate. For pseudotype production, 293T cells were cotransfected with plasmids encoding pNL4-3.Luc.R-E- and VSV-G. The culture medium was exchanged at 6-7 h post transfection, and the cellular supernatant was harvested at 72 h post transfection. The supernatants were clarified from debris by centrifugation (5 min, 3488 x g, RT) filtered through a 0.45 µm filter, aliquoted and stored at -80 °C.

### Amplification of SIV in T cells and macrophages

To produce SIVmac239/316 Env in CD4^+^ T cells, the concanavalin A stimulated cells were infected with SIVmac239/316 Env generated in 293T cells at a multiplicity of infection (MOI) of 0.1 in RPMI 1640 medium supplemented with 20 % FCS and P/S at RT under occasional shaking. Subsequently, the cells were grown in medium supplemented with IL-2 and incubated for 48 h. The cells were then washed twice with 5 ml of culture medium, transferred to a new cell culture flask, and cultured for 2 weeks. Every 2-3 d, the cells were pelleted, the supernatant was harvested and the cells were dissolved in fresh media at a density of 2 x 10^6^ cells/ml. The supernatants were processed as described above for pseudotypes and viral capsid protein concentration determined by a p27-antigen capture enzyme linked immunosorbent assay (ABL), following the manufacturer’s instructions. The p27-antigen-positive supernatants from CD4^+^ T cell cultures obtained from 9 donor animals were pooled to create the stock of CD4^+^ T cell-derived SIVmac239/316 Env, referred to as T-SIV throughout the manuscript. To generate SIVmac239/316 Env in macrophages (M-SIV), the same procedure as described above was followed, except that washing and harvesting of the cells were conducted without detaching the cells from the cell culture flask, and no IL-2 was added to the cell culture medium. The M-SIV virus stock was derived from infected cultures established from eight donor animals. To confirm the absence of mutations in env introduced during virus replication, we isolated RNA from M-SIV and T-SIV using the High Pure Viral RNA Kit (Roche), converted it to cDNA with the Cloned AMV First-Strand cDNA Synthesis Kit (Invitrogen), and then sequenced it after PCR amplification. The viral stocks were further characterized for p27-capsid content, as described above, and for infectious units/ml by titration on C8166 cells as described before (Stahl-Hennig et al. 1999).

### Infectivity assays

For the determination of virus stock infectivity, TZM-bl cells were seeded at a density of 10,000 cells per well in a 96-well cell culture plate and allowed to adhere for 2 hours prior to infection. Infection was carried out using p27-capsid protein or MOI normalized SIVmac239/316 Env in a total volume of 100 µl. After 2 h of spin-oculation (870 x g, RT) (O’Doherty et al. 2000), the infection was allowed to continue for 3-4 h at 37 °C. Thereafter, the infection medium was replaced by 200 µl of fresh culture medium and the cells were cultured for 72 h. Subsequently, the cells were lysed and beta-galactosidase activity in lysates was detected using a commercially available kit (Applied Biosystem) following the manufacturer’s protocol. For lectin inhibition assays, infectivity normalized M-SIV, T-SIV, HIV-1 NL4-3 and env-defective NL4-3 pseudotyped with VSV-G were preincubated with PBS or the lectins UEA (Eylabs), GNA (Sigma), or CV-N (Boyd et al. 1997) for 15 min at 37 °C. Subsequently, the lectin-virus mix was added to TZM-bl cells for infection, and the infection efficiency was determined as described earlier. To assess antibody-mediated neutralization, a similar experimental procedure as the lectin inhibition assay was carried out, except that sera obtained from SIVmac239-infected rhesus macaques were used instead of lectins. Before use, the sera were heat-inactivated for 30 minutes at 56 °C.

### Transmission assays

To model viral transmission, 30,000 Raji, Raji DC-SIGN, Raji DC-SIGNR, or Raji Langerin cells were preincubated for 2-3 h at 37 °C with virus adjusted to ensure equivalent infectivity on the CEMx174 R5 target cell line. Subsequently, the cells underwent two washes with 5 ml PBS each (270 x g, 5 min) to eliminate unbound virus. Following this, the cells were co-cultured with 30,000 CEMx174 R5 target cells in 100 μl of RPMI-1640 medium in a 96-well cell culture plates. After two days, 50 μl of the medium was replaced with fresh media. One day thereafter, the cells were lysed, and luciferase activity was quantified utilizing a commercially available assay kit (Promega). In addition, all cell lines were infected directly without subsequent removal of unbound viruses to control the uniform infectivity of viruses and background signals of transmitter cell lines.

### Western blot

For the analysis of viral particle content, the virus was concentrated through a 20 % sucrose cushion in TNE buffer (0.01 M Tris-HCl pH 7.4, 0.15 M NaCl and 2 mM EDTA in ddH_2_O) using centrifugation. The proteins from the pelleted virions were then separated using SDS-PAGE and subjected to western blot analysis. SIV gp120 was detected using the mouse monoclonal gp120-specific antibody DA6 (Edinger et al. 2000) at a dilution of 1:2,000, while the mouse monoclonal p27-specific antibody 55-2F12 (Higgins et al. 1992) was employed at a dilution of 1:100 for the detection of p27-capsid protein. As a secondary antibody, a horseradish peroxidase-labeled antibody of appropriate species specificity from Dianova was employed at a dilution of 1:5,000. In order to examine the glycosylation of gp120, the concentrated virus was treated with either Endo H or PNGase F from New England Biolabs, for 30 min prior to SDS-PAGE. Signal intensities of western blot bands were quantified using the software ImageJ (Schneider et al. 2012).

### Glycoprofiling by xCGE-LIF

To investigate the *N*-glycosylation of gp120 from M- and T-SIV, sample preparation and analysis were performed as described before (Hennig et al. 2015). Briefly, the virions were concentrated by ultra-centrifugation through a sucrose cushion and the viral proteins were separated by SDS-PAGE. The gp120 protein bands were excised from the Coomassie Blue-stained SDS-polyacrylamide gels, destained, reduced, and alkylated. The attached *N*-glycans were then released by in-gel incubation with PNGase F, and the released *N*-glycans were extracted with water. Next, the *N*-glycans were labeled with 8-aminopyrene-1,3,6-trisulfonic acid (APTS), and any excess label was removed using hydrophilic interaction solid phase extraction. The fluorescently labeled *N*-glycans were separated and analyzed by xCGE-LIF. The glyXtoolCE software (glyXera) was utilized to process the data generated by xCGE-LIF, including the normalization of migration times to an internal standard. This resulted in the creation of *N*-glycan “fingerprints” where the signal intensity in relative fluorescence units (RFU) was plotted on the y-axis against the aligned migration time in aligned migration time units (MTU”) on the x-axis. The high reproducibility of aligned migration times allowed for the comparison of *N*-glycan “fingerprints” between different samples. To elucidate the *N*-glycan structures and annotate the peaks, an in-house *N*-glycan database was used. For quantitative comparison, the relative peak height, which represents the ratio of the peak height to the total height of all peaks, was calculated for each peak and sample.

### LC-MS/MS and automated MS data analysis

After the in-gel release of *N*-glycans by PNGase F treatment proteins were digested with trypsin according to the method outlined by Shevchenko *et al*. (Shevchenko et al. 2006). The procedure involved reducing the proteins with 10 mM DTT (Sigma-Aldrich), followed by carbamidomethylation with 100 mM iodocetamide (Sigma-Aldrich), and subsequent digestion with sequencing-grade trypsin (Promega). To extract the resulting peptides, acetonitrile was used, and the samples were then dried in a vacuum centrifuge before being dissolved in a solution containing 2 % (v/v) acetonitrile and 0.1% (v/v) trifluoroacetic acid (Sigma-Aldrich) for subsequent LC-MS/MS analysis. The analysis was performed using a LTQ-Orbitrap Velos mass spectrometer (Thermo Fisher Scientific) coupled online to a nano-flow ultra-high-pressure liquid chromatography system (RSLC, Thermo Fisher Scientific). Reverse-phase chromatography and mass spectrometry was carried out as described previously (Konze et al. 2014). For data analysis, the MaxQuant proteomics software suite version 1.2.2.5 (Cox & Mann 2008) was utilized, and peak lists were searched against the SIVmac239/316 Env sequence using the Andromeda search engine version 1.1.0.36 (Cox et al. 2011).

### Software

Graphs and statistics were conducted using the GraphPad Prism 9 (Dotmatics) software unless stated otherwise. The text of this manuscript was subjected to rephrasing using ChatGPT (OpenAI) to enhance its linguistic quality.

## Acknowledgements

We extend our gratitude to the following individuals and organizations for their contributions: K. L. Clayton for providing critical feedback on the manuscript, K. Gustafson for providing cyanovirin-N, J. Münch for the plasmids encoding proviral DNA, and R. Desrosiers for antibody DA6. We also acknowledge the NIH AIDS Reagent Program, Division of AIDS, NIAID, NIH, for providing the following reagents: TZM-bl (Cat #8129) from J.C. Kappes, X. Wu, and Tranzyme Inc., B-THP-1 (Raji) and B-THP-1/DC-SIGN (Raji DC-SIGN) from Drs. Li Wu and Vineet N. KewalRaman, human recombinant interleukin-2 (Cat #136) from M. Gately, Hoffmann - La Roche Inc., His-tagged griffithsin (Cat #11610) from B. O’Keefe and J. McMahon, and SIVmac p27 monoclonal antibody 55-2F12 (Cat #1610) from N. Pedersen. Mass spectrometry analysis was carried out at the “Core Unit Mass Spectrometry - Proteomics,” led by A. Pich, at the Institute of Toxicology, Hannover Medical School. The following reagents were obtained through the National Institutes of Health (NIH) AIDS Reagent Program, Division of AIDS, National Institute of Allergy and Infectious Diseases (NIAID), NIH: B-THP-1 and B-THP-1/DC-SIGN from Drs. Li Wu and Vineet N. KewalRamani as well as TZM-bl cells from Dr. John C. Kappes, Dr. Xiaoyun Wu and Tranzyme Inc.

## Abbreviations

AIDS: Acquired immunodeficiency syndrome
APTS: 8-aminopyrene-1,3,6-trisulfonic acid
CBA: Carbohydrate binding agent
CCR5: C-C motif chemokine receptor 5
C.p.s.: Counts per second
CV-N: Cyanovirin-N
CXCR4: C-X-C motif chemokine receptor 4
DC-SIGN: Dendritic-specific intercellular adhesion molecule-grabbing nonintegrin
DC-SIGNR: DC-SIGN related protein
ELISA: Enzyme-linked Immunosorbent assay
Endo H: Endoglycosidase H
Env: Envelope protein
FBS: Fetal bovine serum
FITC: Fluorescein
GNA: Galanthus nivalis agglutinin
HIV: Human immunodeficiency virus
IL-2: Interleukin-2
LC-MS/MS: Liquid chromatography-tandem mass spectrometry
M-SIV: SIV produced in macrophages
MOI: Multiplicity of infection
MTU”: Migration time units
PBMCs: Peripheral blood mononuclear cells
PE-Cy7: Phycoerythrin-cyanine 7
PNGase F: Peptide-N-glycosidase F
P/S: Penicillin/streptomycin
RFU: Relative fluorescence units
RT: Room temperature
RT-PCR: Reverse transcription-polymerase chain reaction
SEM: standard error of the mean
SIV: Simian immunodeficiency virus
tpi: total peak intensity
T-SIV: SIV produced in CD4+ T cells
UEA: Ulex europaeus agglutinin
VSV-G: Vesicular stomatitis virus glycoprotein
wpi: weeks post infection
xCGE-LIF: Multiplexed capillary gel electrophoresis with laser-induced fluorescence detection

## Data Availability statement

The data underlying this article will be shared on reasonable request to the corresponding author.

## Author’s contributions

Conceptualization: R.G.-S. and S.P.; supervision: R.G.-S. and S.P.; project administration: C.B.K, C.S.-H., E.R., and U.R.; resources: R.G.-S., E.R., and C.S.-H.; methodology: C.B.K., F.F.R.B., B.R., A.K., E.R., C.S.-H., and S.P.; investigation: C.B.K., F.F.R.B., S.C., I.N., B.R., A.K., N.S.-L., and C.S.-H.; data curation: C.B.K., S.C., B.R., A.K., U.S., and C.S.-H.; formal analysis: C.B.K., F.F.R.B., S.C., B.R., A.K., U.S., C.S.-H., and S.P.; validation: C.B.K., F.F.R., S.C., U.S., E.R., C.S.-H. and S.P.; visualization: C.B.K., S.C., B.R., A.K., C.S.-H., and S.P.; writing – original draft: C.B.K. and S.P.; writing – review & editing: C.B.K., F.F.R.B., S.C., I.N., B.R., A.K., U.S., U.S., N.S.-L., U.R., R.G.-S., E.R., C.S.-H., and S.P.; funding acquisition: R.G.-S. and S.P.

Supplemental table 1. Structures and relative intensities of *N*-glycans derived from M-SIV and T-SIV, analyzed by xCGE-LIF. Relative peak abundances are presented as percentages of the total peak intensity (peaks 1-96 = 100%). *N*-glycan structures were assigned to peaks based on migration times matching the entries of an in-house *N*-glycan database. Symbols used to depict *N*-glycan structures are given in figure 2A.

---

**Fig. S1:**
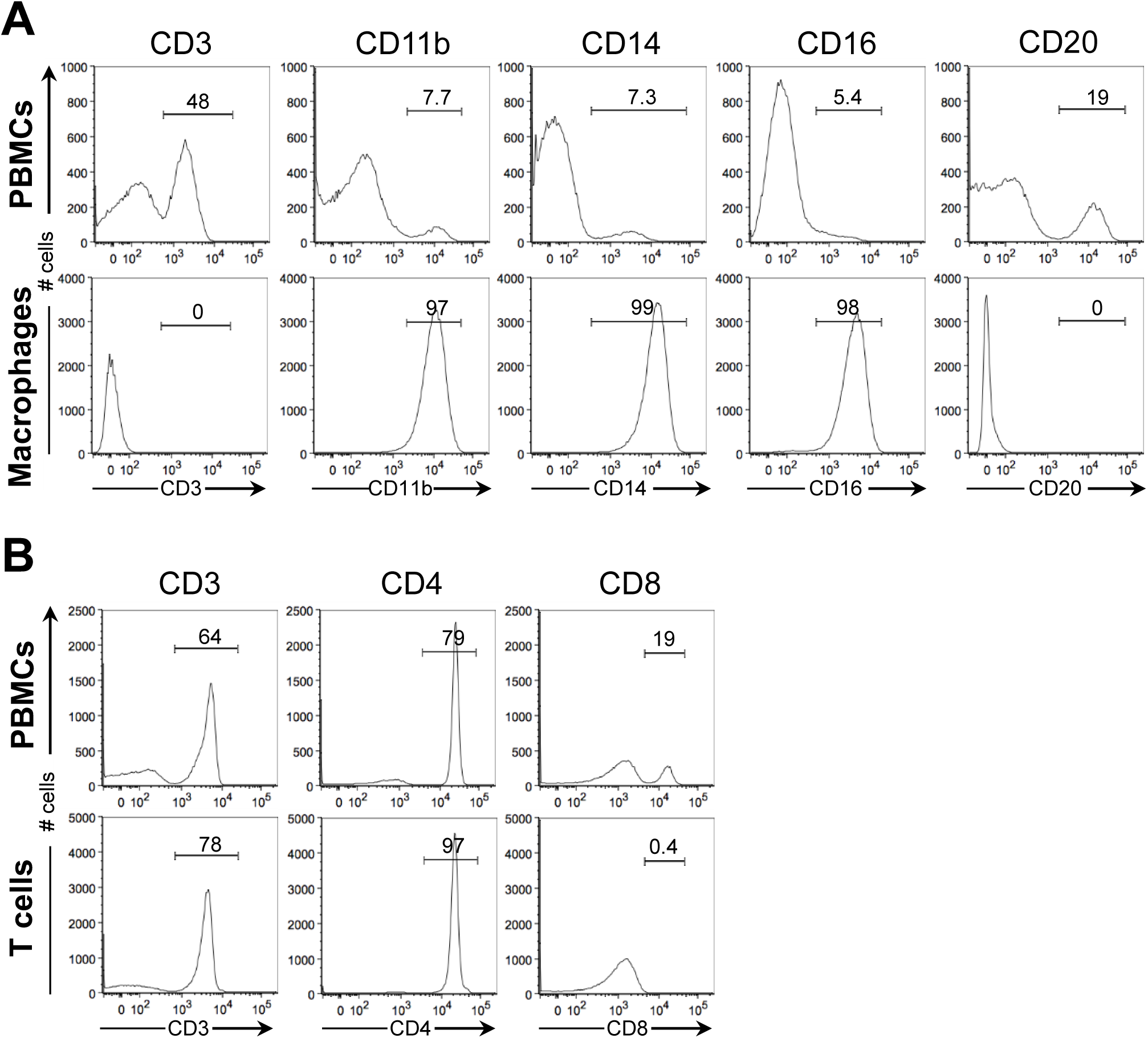
Exemplary flow cytometric validation of purified CD4^+^ T cells and macrophages. A) Rhesus macaque PBMCs and monocyte-derived macrophages were flow cytometrically stained using antibodies targeting macrophage (CD11b, CD14, CD16), T cell (CD3), and B cell (CD20) markers. Representative data from four different experiments are presented. B) Rhesus macaque PBMCs or purified CD4^+^ T cells were stained for flow cytometry using antibodies specific for T cells (CD3) or T cell subpopulations (CD4, CD8). Representative data from two independent experiments are shown. For A) and B), the y-axis represents the cell count, while the x-axis indicates marker signal intensity. Proportions of cells within gates are denoted above the bars.

**Fig. S2:**
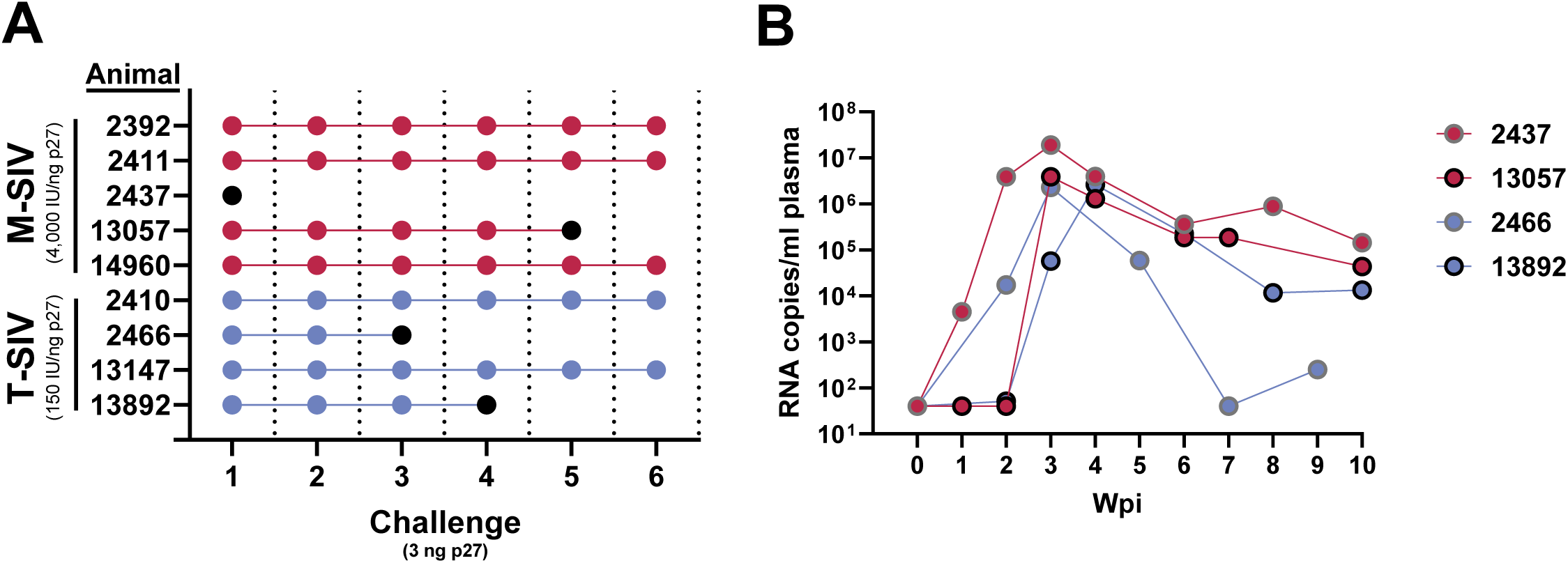
Both M-SIV and T-SIV are infectious *in vivo*. A) Rhesus macaques (n = 4-5 per group) were rectally challenged with 3 ng p27-capsid-protein of M-SIV or T-SIV diluted in PBS. The challenges were repeated every three weeks until the animals became infected (indicated by black filled symbols) or up to six challenges. Infection was determined by detection of SIV RNA in the peripheral blood by quantitative reverse transcriptase-polymerase chain reaction (RT-PCR). Animal identifiers are indicated on the y-axis. B) Plasma viral load of rhesus macaques infected with M-SIV or T-SIV was measured as RNA copies/ml from the day of challenge (day 0) up to 10 weeks post infection (wpi). At 3 wpi with SIVmac239/316 Env, animal 13057 underwent an additional challenge with SIVmac251 as part of a separate experiment, which was not part of this study.

