## Supplemental Table 1 for "Macrophage- and CD4^+^ T cell-derived SIV differ in glycosylation, infectivity and neutralization sensitivity"

| TABLE S1. Structures and relative intensities of N-glycans derived from M-SIV and T-SIV analyzed by xCGE-LIF | | | | | |
|---|---|---|---|---|---|
| N-glycan structure | % Intensity | | N-glycan structure | % Intensity | |
| | M-SIV | T-SIV | | M-SIV | T-SIV |
| 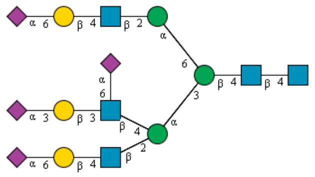 | 1.17 | 2.77 | 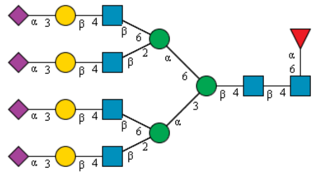 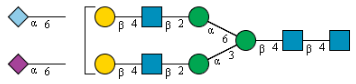 | 0.33 | 1.62 |
| 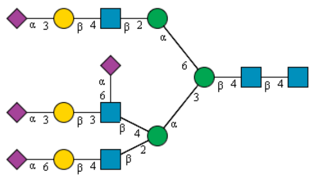 | 1.53 | 3.10 | 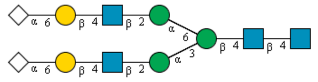 | 0.62 | 1.72 |
| 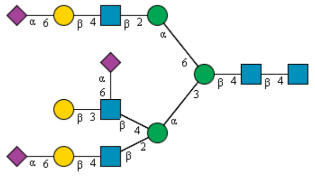 | 1.46 | 1.33 | 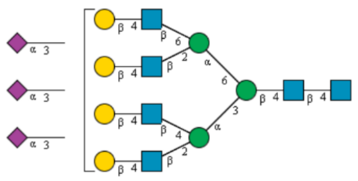 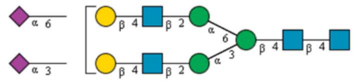 | 3.15 | 2.39 |
| 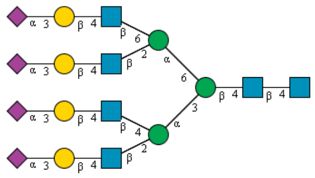 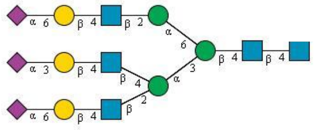 | 1.78 | 2.54 | 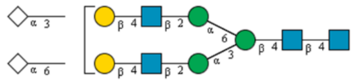 | 3.22 | 3.31 |
| 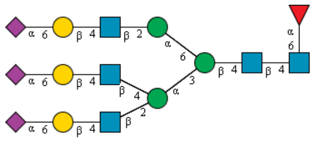 | 2.72 | 4.32 | 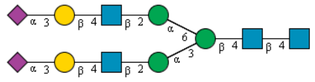 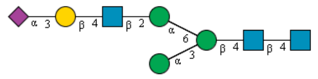 | 1.28 | 0.84 |
| | | | 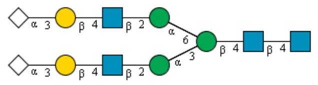 | 0.00 | 7.97 |
| | | | 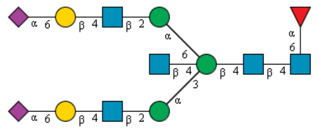 | 1.79 | 0.00 |

| | | | | | |
|---|---|---|---|---|---|
| | | | 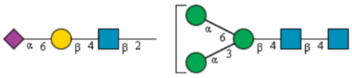 | | |
| 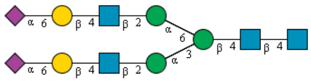 | continued | | 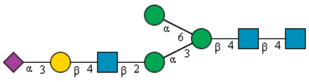 | 2.42 | 1.76 |
| 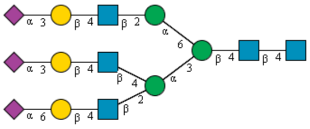 | | | 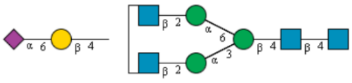 | 0.30 | 0.00 |

TABLE S1. (continued)

| N-glycan structure | % Intensity | | N-glycan structure | % Intensity | |
|---|---|---|---|---|---|
| | M-SIV | T-SIV | | M-SIV | T-SIV |
| 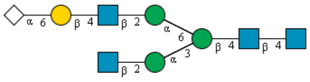 | 0.00 | 0.55 | 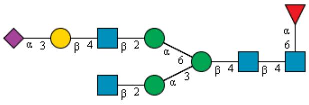 | 0.75 | 1.34 |
| 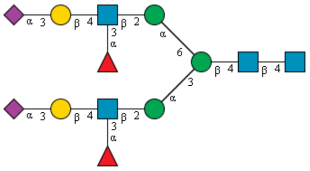 | | | 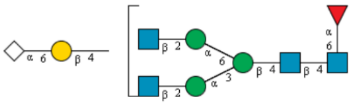 | | |
| 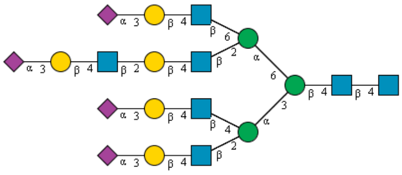 | 1.72 | 1.35 | 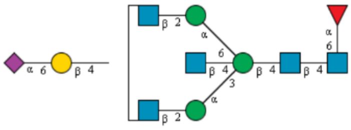 | | |
| 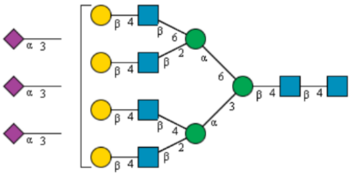 | | | 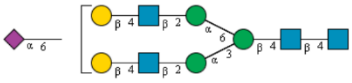 | 1.20 | 0.00 |
| | | | 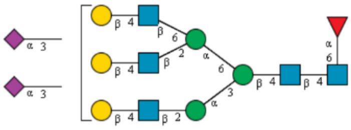 | 0.00 | 0.93 |
| 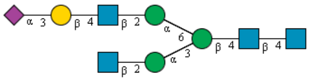 | 2.37 | 1.02 | 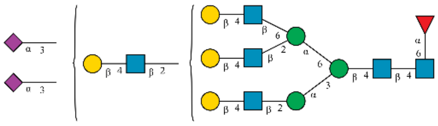 | | |

| N-glycan structure | M-SIV | T-SIV | N-glycan structure | M-SIV | T-SIV |
|---|---|---|---|---|---|
| 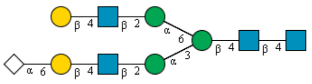 | 0.00 | 1.45 | 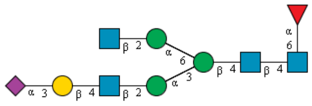 | 0.15 | 0.00 |
| 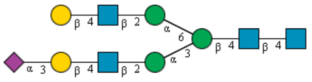 | 4.61 | 0.00 | 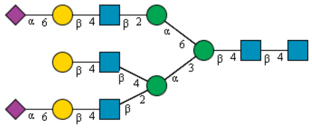 | 0.54 | 0.71 |

TABLE S1. (continued)

| N-glycan structure | % Intensity | | N-glycan structure | % Intensity | |
|---|---|---|---|---|---|
| | M-SIV | T-SIV | | M-SIV | T-SIV |
| 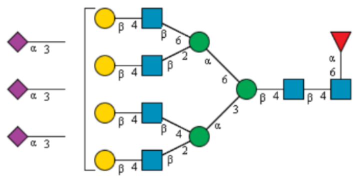 | 1.30 | 1.55 | 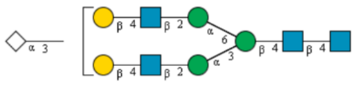 | 0.00 | 2.03 |
| 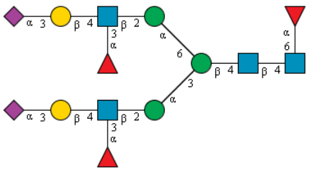 | | | 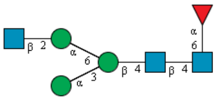 | 3.86 | 0.00 |
| 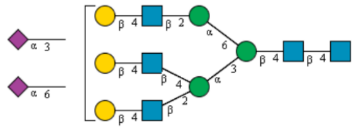 | | | 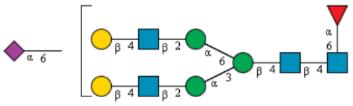 | | |
| 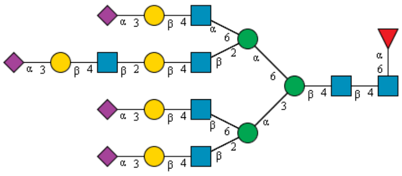 | | | 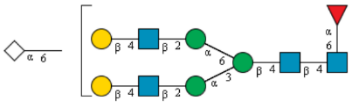 | | |
| 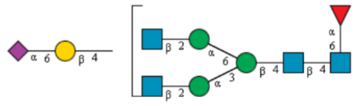 | 2.07 | 2.24 | 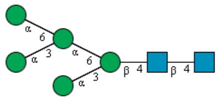 | 2.28 | 7.17 |
| 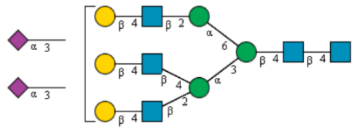 | | | 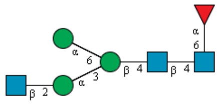 | | |

| | | | | | |
|---|---|---|---|---|---|
| 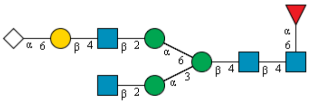 | | | 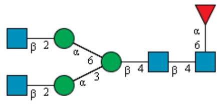 | 0.10 | 0.11 |
| 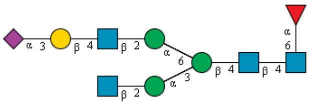 | 0.75 | 1.34 | 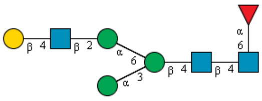 | 0.88 | 1.37 |
| 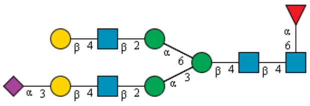 | | | 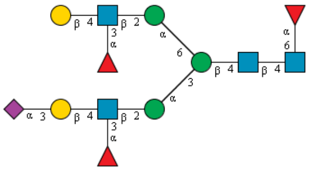 | | continued |
| 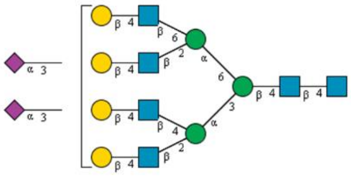 | 1.50 | 1.15 | 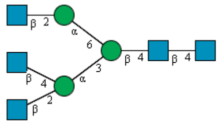 | 1.49 | 0.55 |
| 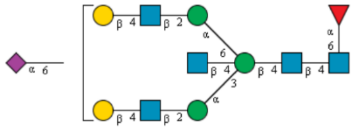 | | | 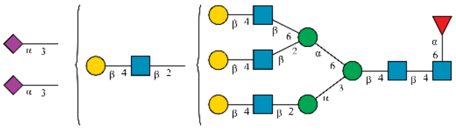 | | |

TABLE S1. (continued)

| N-glycan structure | % Intensity | | N-glycan structure | % Intensity | |
|---|---|---|---|---|---|
| | M-SIV | T-SIV | | M-SIV | T-SIV |
| 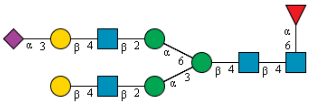 | | | 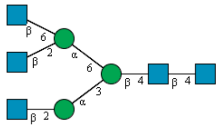 | 0.27 | 0.00 |
| 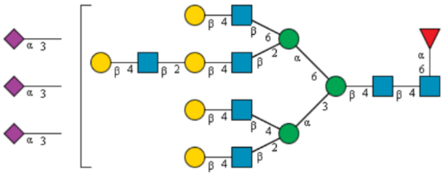 | 4.54 | 1.73 | 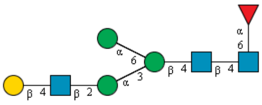 | 0.32 | 2.86 |
| 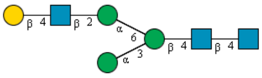 | 1.38 | 0.72 | 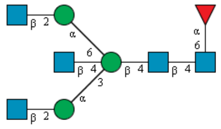 | 0.25 | 0.32 |

| | | | | | |
|---|---|---|---|---|---|
| 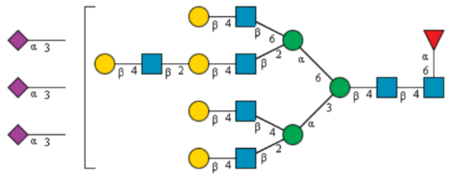 | 1.76 | 0.50 | 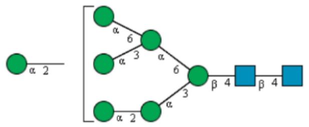 | 1.49 | 0.48 |
| 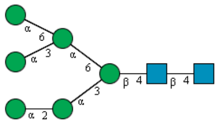 | 0.75 | 1.79 | 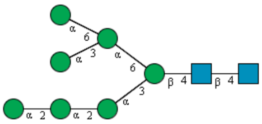 | 1.41 | 0.72 |
| 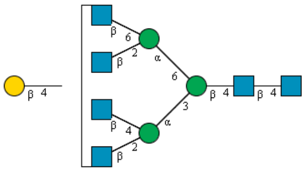 | 2.16 | 1.37 | 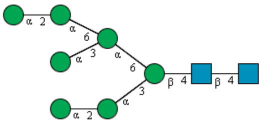 | 0.60 | 0.71 |

TABLE S1. (continued)

| N-glycan structure | % Intensity | | N-glycan structure | % Intensity | |
|---|---|---|---|---|---|
| | M-SIV | T-SIV | | M-SIV | T-SIV |
| 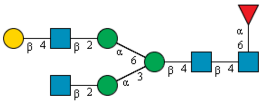 | 0.37 | 0.56 | 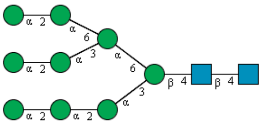 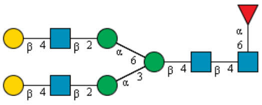 | 0.98 | 1.93 |
| 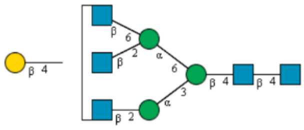 | 0.60 | 0.14 | | | |
| 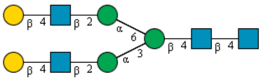 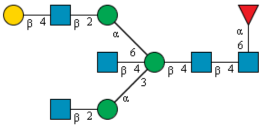 | 0.30 | 0.31 | 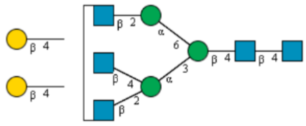 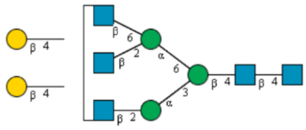 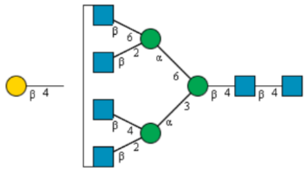 | 2.02 | 0.92 |
| 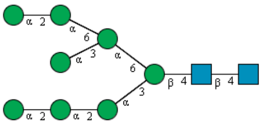 | 0.61 | 3.34 | | | |

| | | | | | |
|---|---|---|---|---|---|
| 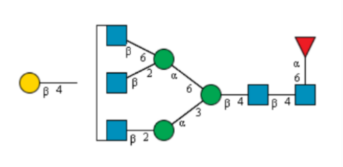 | 0.25 | 0.26 | 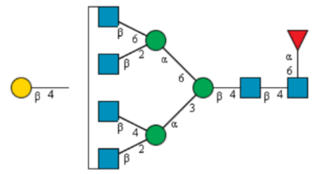 | 0.26 | 0.40 |
| 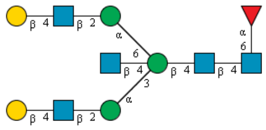 | 0.00 | 0.18 | 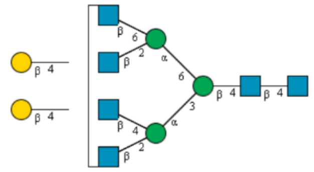 | 0.76 | 0.38 |
| 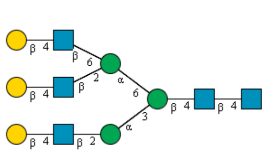 | 0.19 | 0.00 | 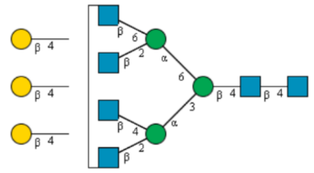 | 0.55 | 0.16 |

TABLE S1. (continued)

| N-glycan structure | % Intensity | | N-glycan structure | % Intensity | |
|---|---|---|---|---|---|
| | M-SIV | T-SIV | | M-SIV | T-SIV |
| 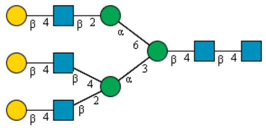 | 0.66 | 0.39 | 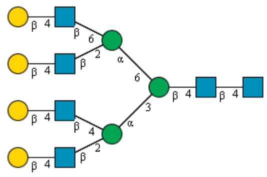 | 0.27 | 0.14 |
| 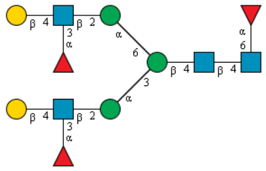 | 0.75 | 0.70 | 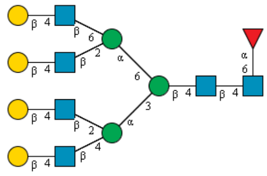 | 0.15 | 0.29 |
| 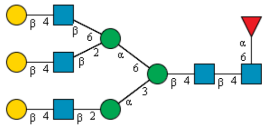 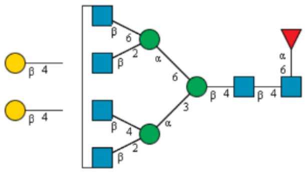 | 0.00 | 0.29 | 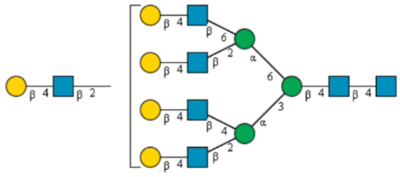 | 0.96 | 0.75 |
